# Locoregional Radiogenomic Models Capture Gene Expression Heterogeneity in Glioblastoma

**DOI:** 10.1101/304105

**Authors:** Adrien Depeursinge, Tünde Szilágyi, Yan Liu, Kázmèr Kovács, Reena P. Thomas, Kristen W. Yeom, Nancy Fischbein, Daniel L. Rubin, Michael, Olivier Gevaert

**Affiliations:** Biomedical Imaging Group, École polytechnique fédérale de Lausanne (EPFL), Lausanne 1015, Switzerland; Institute of Information Systems, University of Applied Sciences Western Switzerland (HES-SO), Sierre 3960, Switzerland; Stanford Center for Biomedical Informatics Research, Department of Medicine, Stanford University, Stanford CA, USA; Department of Radiology, the Second Affiliated Hospital, Zhejiang University School of Medicine, Hangzhou, China; Department of Medical Imaging, Faculty of Medicine, University of Debrecen, Debrecen, Hungary; Department of Neurology, Stanford University, Stanford CA, USA; Department of Radiology, Stanford University, Stanford CA, USA; Department of Biomedical Data Science, Stanford University, Stanford CA, USA

## Abstract

Radiogenomics mapping noninvasively determines important relationships between the molecular genotype and imaging phenotype of various tumors, allowing advances in both clinical care and cancer research. While early work has shown its technical feasibility, here we extend radiogenomic mapping to a locoregional level that can account for the molecular heterogeneity of tumors. To achieve this, our data processing pipeline relies on three main steps: 1) the use of multi-omics data fusion to generate a set of 100 interpretable gene modules, 2) the use of patch-based image analysis (specifically of contrast-enhanced T1-weighted weighted MR images) combined with Generalized Linear Models (GLM) to establish potential links between module expressions and local MR signal, and 3) the use of expression heatmaps based on GLMs decision values to explore visualization of tumor molecular heterogeneity. The performance of the proposed approach was evaluated using a leave-one-patient-out crossvalidation method as well as a separate validation data set. The top performing models were based on a small set of 20 features and yielded Area Under the receiver operating characteristic Curve (AUC) above 0.65 on the validation cohort for eight modules. Next, we demonstrate the clinical and biological interpretation of four modules using molecular expression heatmaps superimposed on clinical radiographic images, showing the potential for assessing tumor molecular heterogeneity and the utility of this method for precision treatment in clinical decision making and imaging surveillance.

## Introduction

Radiogenomics is an burgeoning field of science that reveals important relationships between molecular properties of tumors and their macroscopic appearance on radiographic imaging used in the clinical setting. It is defined by the linking of quantitative image features with molecular data and holds the potential to be readily translatable to clinical decision making by the identification of imaging biomarkers that can accurately predict underlying tumor biology and the efficacy of treatment. The actual translation of this approach to the clinic is feasible as medical imaging is a standard part of the routine diagnostic work-up and surveillance of cancer patients.

It is well known that there is genomic heterogeneity between tumors and also within individual tumors ^1, 2^. However, the only way to determine tumor genotype is through invasive surgical sampling, and given the spatial inhomogeneity of genomic expression within a tumor, information about the local *in vivo* microenvironment^3^ and locoregional tumor precursor cells-of-origin ^4^ can be lost in biopsies. The creation of radiogenomic maps bypasses the need for surgical sampling in order to understand a tumor lesions’ underlying biology and the changes that may result from treatment intervention. The application of radiogenomic maps has been demonstrated in hepatocellular carcinoma ^5^, lung cancer ^6–8^, and gliomas (particularly high grade gliomas such as glioblastoma) ^9–12^. The aforementioned analyses utilized human-based semantic descriptions of the tumor appearance on imaging such as the Visually AcceSsible Rembrandt Images (VASARI) features ^13^, while others utilized global imaging features such as the extent of tumor associated vasogenic edema, regions of non-enhancing tumor, enhancement and necrosis from semi-automatic segmentations of tumors with imaging analysis software ^12^, and lastly, global extraction of more classical radiomics quantitative features extracted across the gross tumor volume ^14^. In contrast, radiogenomics models in the two latter studies demonstrate significant but average performance. Importantly, there exists a significant body of literature confirming that molecular expression is variable within a single tumor lesion and, therefore, studies which assume homogeneity across the entire volume of tumor may lead to an oversimplified and inaccurate radiogenomics model. To this end, recent studies investigated a locoregional approach based on manually delineated necrotic, enhancing, edematous or nonenhancing tumor regions ^15,16^. Beig *et al.* ^15^ observed that the localized radiomics features that were most informative of hypoxia distinguished between short- and long-term survivors. Similarly, Hu *et al.* ^16^ revealed significant imaging correlations with local expression of molecular markers in glioblastoma. This study had a low sample size of only 13 tumors due to the need for multiple localized image-guided biopsies and the unique challenge in the clinical setting of brain tumors requiring neurosurgical intervention which may not be clinically indicated.

In this study, we implement radiogenomics mapping that accounts for the locoregional heterogeneity of brain tumors as observed on imaging and link signature image features with their molecular profiles. We use an automated and localized image analysis by dividing the tumors into patches. The latter is based on the hypothesis that confined tumor habitats have distinct molecular and imaging signatures when compared to larger tumor volumes ^17^. Moreover, this provides the opportunity to estimate local molecular profiles and further relate them using a holistic approach to assess brain tumors. We focus this study on human glioblastoma (GBM), which is the most common and most lethal form of primary brain cancer in adults, accounting for 16% of all primary tumors and 82% of all malignant tumors within the central nervous system ^18^. GBM occurs most frequently within the 45-70 year age group with a mean age of 63 years in the USA, and is 1.6 times more prevalent in males than females ^18^. GBMs are typically fast-growing and infiltrative tumors that are invariably fatal. With current standardized treatment, the median survival rate for patients with glioblastoma is just 14.6 months, and the five-year overall survival rate is 4.7% ^18^.

Our work shows that locoregional modeling of Magnetic Resonance (MR) images of GBM can capture gene module expression and heterogeneity within this tumor.

## Methods

### Study populations multi-omics and MR imaging data

We used two distinct and publicly available radiogenomics datasets: we first used patients from The Cancer Genome Atlas (TCGA) glioblastoma data set^19^ to develop locoregional radiogenomics maps, and then validated them using the REpository of Molecular BRAin Neoplasia DaTa (REMBRANDT) dataset ^20,21^. We used multi-omics molecular data from TCGA project including gene expression, DNA methylation and DNA copy number data ^19^. The gene expression data were produced using Agilent microarrays for GBM. Preprocessing was done by log-transformation and quantile normalization of the arrays ^22,23^. The DNA methylation data were generated using the Illumina Infinium Human Methylation 27 Bead Chip. DNA methylation was quantified using β-values ranging from 0 to 1 according to the DNA methylation levels. We removed CpG sites with more than 10% of missing values in all samples. We used the 15-K nearest neighbor algorithm to estimate the remaining missing values in the data set^24^. Finally, the copy number data we used were produced by the Agilent Sure Print G3 Human CGH Microarray Kit 1M× 1M platform. This platform has high redundancy at the gene level, but we observed high correlation between probes matching the same gene. Therefore, probes matching the same gene were merged by taking the average. For all data sources, gene annotation was translated to official gene symbols based on the HUGO Gene Nomenclature Committee identifiers. Next, TCGA samples were analyzed in batches and a significant batch effect was observed based on a one-way analysis of variance in most data modes. We applied Combat to adjust for these effects ^25^. For the REMBRANT cohort only gene expression data was available produced by Affymetrix GeneChip Human Genome U133 Plus 2.0 microarrays archived by G-DOC ^26^. Preprocessing was done as previously reported ^27^. Briefly the RMA algorithm was used for preprocessing the microarray data ^28^. Matched Magnetic Resonance (MR) images were available for both the TCGA and REMBRANDT cohorts from The Cancer Imaging Archive (TCIA) ^29^. We used only pre-surgical, gadolinium-enhanced, axial T1-weighted images with a slice thickness equal to or less than 3 mm. This resulted in a total of 84 and 47 patients for the TCGA and REMBRANDT cohorts, respectively.

### Multi-omics data fusion

We used AMARETTO, a multi-omics data fusion framework creating gene expression modules by integrating DNA methylation, DNA copy number and gene expression data ^30–32^ using the TCGA data and created the gene expression modules on the REMBRANDT data set. To generate a gene regulatory module network of glioma, we first applied AMARETTO on the preprocessed copy number, DNA methylation and gene expression data. DNA methylation data was modeled using MethylMix ^33–37^ identifying only differentially methylated and transcriptionally predictive genes. Genomic Identification of Significant Targets in Cancer (GISTIC) was used to identify the recurrently amplified and deleted genes using DNA copy number data. Both lists of genes were used in combination with RNA gene expression data to develop 100 AMARETTO gene modules ^30,31,38^ linking the genomic- and epigenomics-driven genes to their targets. Modules were annotated using gene set enrichment analysis. We used the following gene-set databases: MSigDB version 3 ^39^, GeneSetDB version 4 ^40^, CHEA for CHIP-X gene sets version 2 ^41^ and manually curated gene sets related to stem cells and immune gene sets. We use a hypergeometric test to check for enrichment of gene sets in the lists of hyper- and hypo-methylated genes. We corrected for multiple testing using the False Discovery Rate (FDR) ^42^. Modules were then created for the REMBRANDT data set using the regulator and target gene expression values. Modules were represented by the mean average expression of all target genes in each module. Note that only gene expression data is necessary to represent the modules, and no DNA methylation or DNA copy number data is needed. All module values were standardized to have mean zero and standard deviation equal to one.

### MR Image analysis

Images from the TCGA and REMBRANDT cohorts were processed in the following manner: Regions Of Interests (ROIs) encompassing the enhancing tumor area on all contiguous axial slices were outlined by a panel of experienced radiologists and neuroradiologists by consensus (KY, YL and MI). The pre-contrast T1 images were reviewed to ensure ROI performance strictly on the enhancing component. Bias correction was applied to all images using the FSL 5.0.9 FAST algorithm ^43^. All axial slices from all image series were resampled using bi-cubic interpolation to have a pixel size of 0.5 ×0.5 mm^2^. Voxel values were standardized to have an average of zero and unit variance inside the brain. The latter was achieved by fitting a mixture of two Gaussians parameterized by (*μ*_0_,*σ*_0_) and (μ_1_,*σ*_1_) using an iterative Expectation-Maximization algorithm and imposing *μ*_1_ — *μ*_0_ ≥ 50. The Gaussian model (*μ*_1;_ *σ*_1_) is capturing extended brain values. Standardized MR data was obtained by using the brain mean value *μ*_1_ and standard deviation value σ_1_. The REMBRANDT preprocessed data and segmentations are made publicly available on the TCIA wiki (doi: 10.7937/K9/TCIA.2018.3v6dl662).

Next, we divided the tumor region into patches according to the following procedure (Figure 1). The ROIs were divided into overlapping circular patches with a radius of *r*=12 pixels (6mm) in axial slices and center *x*. *r* = 6mm was chosen as a trade-off between locality and the wealth of visual information encompassed in the patches ^44^. The patches were randomly positioned while keeping a minimum distance between patch centers equal to their radius *r*. The latter process was repeated 50 times to investigate robustness to patch positioning. Two sets of patches were extracted: from the boundary *b* and from the core c of the ROI. The boundary patches were centered on the tumor contour (Figure 1) and restricted to keep a minimum distance *r* between their centers. The core patches can lie anywhere inside the ROI while keeping a minimum distance r between both their centers as well as the tumor contour (Figure 1). The four lower- and uppermost axial slices containing the tumor were discarded to avoid partial volume effects. An average of 23,981 boundary and 21,129 core patches were obtained from a total of 1,527 slices for the 84 TCGA patients. Similarly, 7,894 boundary and 8,004 core patches were obtained from 425 slices for REMBRANDT patients. For each patient *p*, boundary and core feature vectors *η_i_ =* (*η_i,m=1,…,20_*) were extracted to construct the feature matrices from the collections of *K* boundary and *L* core patches centered at locations 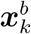 and 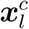, respectively. Twenty intensity features were obtained per patch as the counts in each histogram bins in the interval [-1,1] with a bin width of 0.1. Boundary and core texture features were obtained from responses of Circular Harmonic Wavelets (CHW), which consists of a filterbank analyzing local circular frequencies at multiple scales ^45,46^. The responses of the filters aggregated over the boundary patches provide tumor margin descriptors, whereas their responses aggregated over the core patches yield texture attributes ^47^. The wavelet coefficients (i.e., the responses of the filters) were obtained by convolving the MRI slice with every filter, and the mean energies of the coefficients were computed over an image patch to provide scalar feature values. Three dyadic scales were used to cover the circular patches. Circular harmonic sets in [0, *N*] were tested with *N* = 0,…, 5, yielding feature vectors of size 3(*N* + 1).

**Figure 1.**
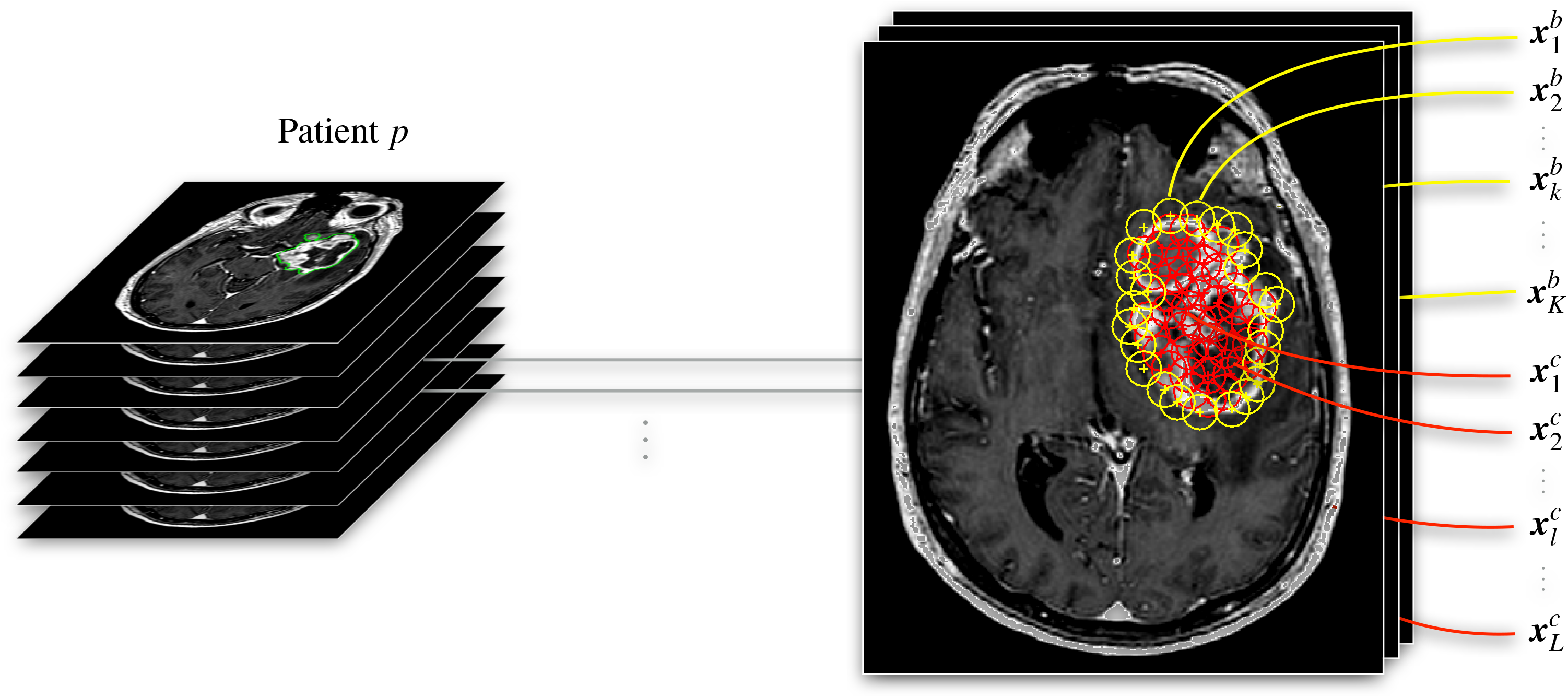
Workflow for radiogenomic modeling using patches by dividing the tumor region into overlapping circular patches with a radius of 6mm. We used two types of patches, core and boundary patches, shown in red and yellow respectively.

### Radiogenomics modeling

The over- and under-expression of the gene modules was predicted on a patch basis using a multivariate linear model based on the vectors of intensity features *η_i_*. One fourth of the patients with most extreme gene expression were chosen in order to define sets of exemplar patients for each gene module *q*. Therefore, among the 84 and 47 patients of TCGA and REMBRANDT respectively, 10 and 5 patients with most negative module expression and 10 and 5 patients with most positive module expression were selected per gene module to represent it. Three validation strategies were used to estimate the predictive performance of each module-specific model: (1) a Leave-One-Patient-Out Cross Validation (LOPO-CV) on the 20 extreme patients selected per module from the TCGA dataset, (2) training the model with TCGA and validating it on REMBRANDT, and (3) a LOPO-CV on the 10 extreme patients selected per module from the REMBRANDT dataset. When using LOPO-CV, we carefully separated patches by patients between the training and test sets. Receiver Operating Characteristic (ROC) analysis was carried out on a patch basis. Separate models were built for boundary and core patches. The class membership *y_q,i_* (i.e., over-versus under-expressed gene module *q*) of the patch *i* expressed in terms of its feature vector *η_i_* was estimated using a Generalized Linear Model (GLM) ^48^ as

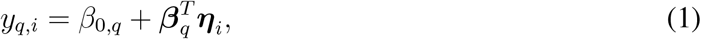

where the decision value *y_q,i_* = [-1,1] for under- and over-expression of *q* in a training patch *i*.

## Results

### AMARETTO modules capture major glioblastoma pathways

We used AMARETTO to build 100 gene expression modules based on 426 TCGA patients. Each module was further annotated using gene set enrichment analysis (Supplementary Table 1) resulting in modules enriched in key biological pathways including apoptosis, cell cycle, immune response and angiogenesis (Figure 2). Next, we validated these modules using the gene expression data of the REMBRANDT validation cohort. Approximately two thirds of the modules have high R-square and module homogeneity in the REMBRANDT cohort (Supplementary Table 2).

**Figure 2.**
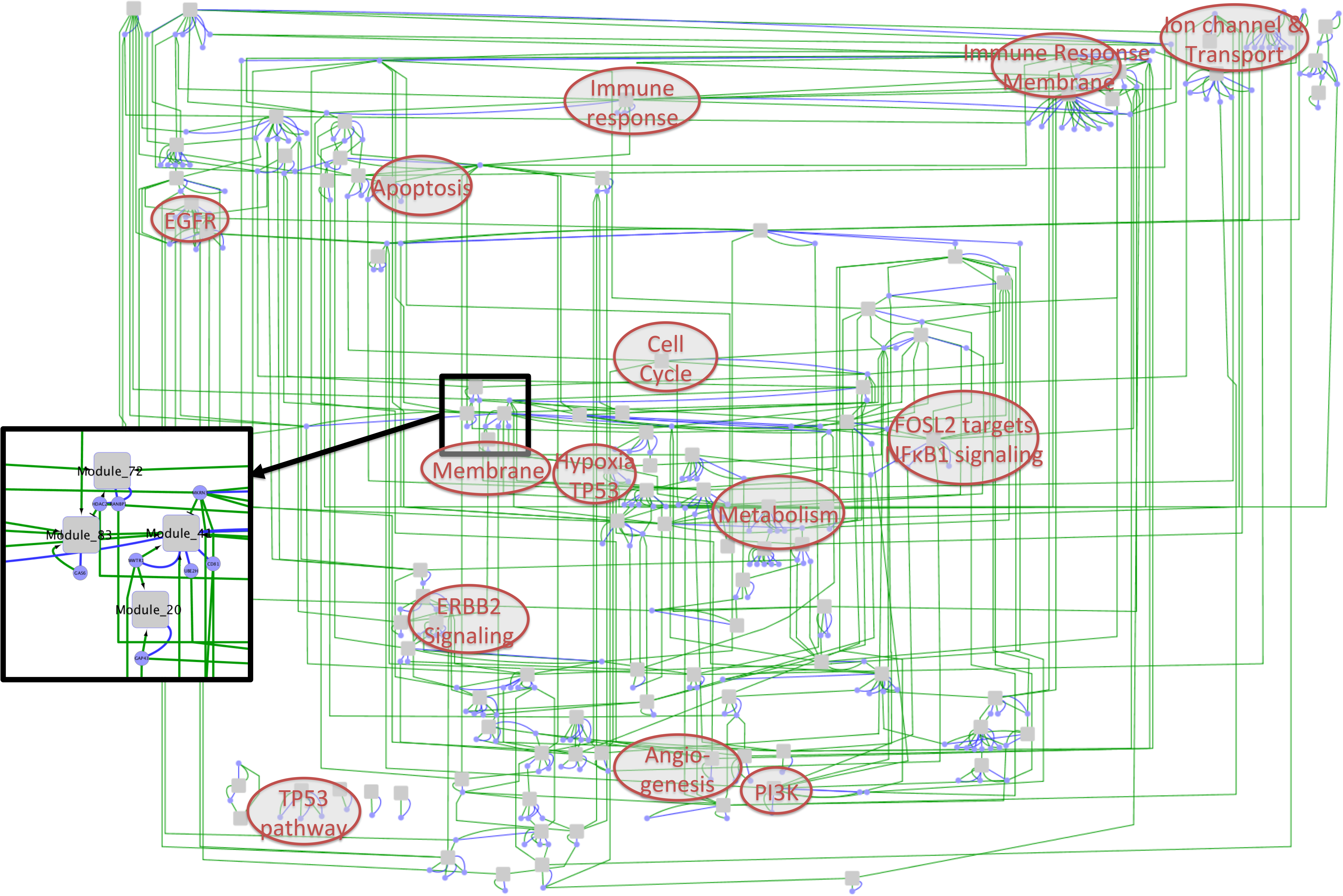
Visualization of the relationships between the gene expression modules. Grey nodes are modules (co-expressed gene sets); blue nodes are cancer driver genes. Blue links represent membership of a cancer driver to a module, and green lines represent membership to a regulatory program of a module. We have annotated key gene set enrichments on the modules. A magnified part of the network is displayed on the left showing four modules including Module 20 together with their cancer driver genes.

### Radiogenomics modeling

For each module the Area Under the ROC Curve (AUC) was measured based on the GLM’s continuous decision values. AUCs were computed for predictions based on either core or boundary patches (Figure 1 and Table 1). AUCs were averaged over 50 repetitions of the random patch positioning. Performance comparison and combination of intensity and texture features show that texture-based models cannot accurately predict the presence or absence of gene modules, and that combining them with intensity harms the predictive performance in most cases (Supplementary Table 3). Therefore, we focused on intensity-based models. AUCs above 0.65 for all three validation setups were observed for modules 70 and 90 when the model was based on core patches as well as for modules 36 and 84 when based on boundary patches. AUCs above 0.65 with both TCGA LOPO and REMBRANDT validation were observed for modules 32, 24 and 45 when the model was based on core patches as well as for module 45 when based on boundary patches. Note that the latter was best predicted using core patches. AUCs above 0.70 with both TCGA LOPO and REMBRANDT LOPO were observed for modules 11, 93, 68, 22, 42, 55, 61 and when based on core patches.

**Table 1.** AUCs associated with radiogenomics modeling. The results are sorted based on TCGA LOPO. Average and standard deviations over the 50 MC repetitions are reported. AUC=Area under the Receiver Operating Characteristic curve, TCGA=The Cancer Genome Atlas, LOPO=Leave-One-Patient-Out cross validation.

| Modules | TCGA | REMBRANDT | REMBRANDT |
| --- | --- | --- | --- |
|  | LOPO | trained on | LOPO |
|  |  | TCGA |  |
|  | Core patches |  |  |
| Module 32 | 0.8786 ± 0.0024 | 0.6569 ± 0.0334 | 0.4826 ± 0.0195 |
| Module 11 | 0.8755 ± 0.0017 | 0.1844 ± 0.0094 | 0.8013 ± 0.0099 |
| Module 93 | 0.8301 ± 0.0029 | 0.3798 ± 0.0104 | 0.8309 ± 0.0054 |
| Module 68 | 0.7917 ± 0.0027 | 0.263 ± 0.0066 | 0.6521 ± 0.0064 |
| Module 24 | 0.7874 ± 0.0061 | 0.7902 ± 0.0065 | 0.5749 ± 0.012 |
| Module 45 | 0.7873 ± 0.0057 | 0.7733 ± 0.006 | 0.5689 ± 0.0112 |
| Module 22 | 0.7667 ± 0.006 | 0.3066 ± 0.0178 | 0.6686 ± 0.0117 |
| Module 42 | 0.7276 ± 0.017 | 0.2099 ± 0.0081 | 0.8038 ± 0.0092 |
| Module 55 | 0.7265 ± 0.0044 | 0.5429 ± 0.0063 | 0.6933 ± 0.0076 |
| Module 90 | 0.7176 ± 0.0089 | 0.6685 ± 0.0277 | 0.6514 ± 0.0064 |
| Module 61 | 0.7006 ± 0.0049 | 0.2576 ± 0.013 | 0.7316 ± 0.01 |
| Module 70 | 0.6827 ± 0.004 | 0.6563 ± 0.0074 | 0.729 ± 0.0072 |
|  | Boundary patches |  |  |
| Module 45 | 0.6632 ± 0.003 | 0.692 ± 0.005 | 0.6327 ± 0.0059 |
| Module 36 | 0.6581 ± 0.0044 | 0.6524 ± 0.0117 | 0.7112 ± 0.0059 |
| Module 84 | 0.6573 ± 0.0028 | 0.7036 ± 0.0118 | 0.7352 ± 0.0082 |

### Biological interpretation

For the core patches, the best results were observed for several modules including modules 32, 11, 93 and 45. Module 32 is enriched in gene sets related to neural differentiation of embryonic stem cells ^49^ and contains genes involved in synaptogenesis, synaptic transmission and transmission of nerve impulse. This module is regulated by QKI, a gene involved in myelination and oligodendrocyte differentiation and QKI has been shown to maintain stemness of glioma stem cells^50^. Module 11 is a core cell cycle module and is regulated by TNFRSF1A and MYCN, among others, both widely documented to be involved in cell cycle progression in many cancers. Module 93 is enriched in genes located in the membrane, receptor tyrosine kinase pathways, and the ERBB pathway supported also by ERBB3 as one of the regulators of Module 93. Module 45 is enriched in processes related to ion transport and is only regulated by one gene, the transmembrane gene FAM57B. Module 24 has few enrichments and corresponds to unknown biology. Next, two modules has good performance across all three test scenarios, modules 90 and 70. Module 90 includes ERBB3 as driver gene and is involved in transmembrane tyrosine kinase activity. Module 70 is enriched in processes related to packaging of telomere ends and RNA polymerase transcription, likely driven by module 70 regulator gene REV3L. For the boundary patches, fewer modules had acceptable performance, including modules 36, 84 and 45, although module 45 is better predicted using core patches. Module 36 is enriched in the TP53 pathway, contains MDM2 and CDK4, and reflects DNA repair processes. Module 84 is enriched in multiple pathways related to development and has TPBG, BCAN, TCF12 and ANXA11 as regulators. This module is supported by the function of BCAN, a gene highly expressed in gliomas and promoting the growth and cell motility of brain tumor cells.

### Visualization of locoregional module expression

The approach used for patch-based module prediction provides the opportunity to reveal and visualize local tumor patterns related to the over- or under-expression of the corresponding gene module on test patients. For each module *q*, heatmaps were generated based on the value *y_q,i_* of the decision function (Equation 1) of GLMs trained on TCGA for every local patch *i* of the REMBRANDT dataset. The maximum absolute value of the score over the tumor slice is normalized to one. Whereas many heatmaps were homogeneous over the tumors, we observed several heatmaps suggesting molecular heterogeneity. For example, heatmaps of Modules 32 and 45 show intriguing heterogeneity across the tumor ROI in the T1 post-gadolinium image of the same patient (Figure 3). Next, the heatmap for module 70 shows heterogeneity across GBM regions highlighting the common patterns across subjects suggestive of over- and under-expression of the gene module (Figure 4). Module 70 is significantly enriched in processes involved in key pathways associated with RNA transcription.

**Figure 3.**
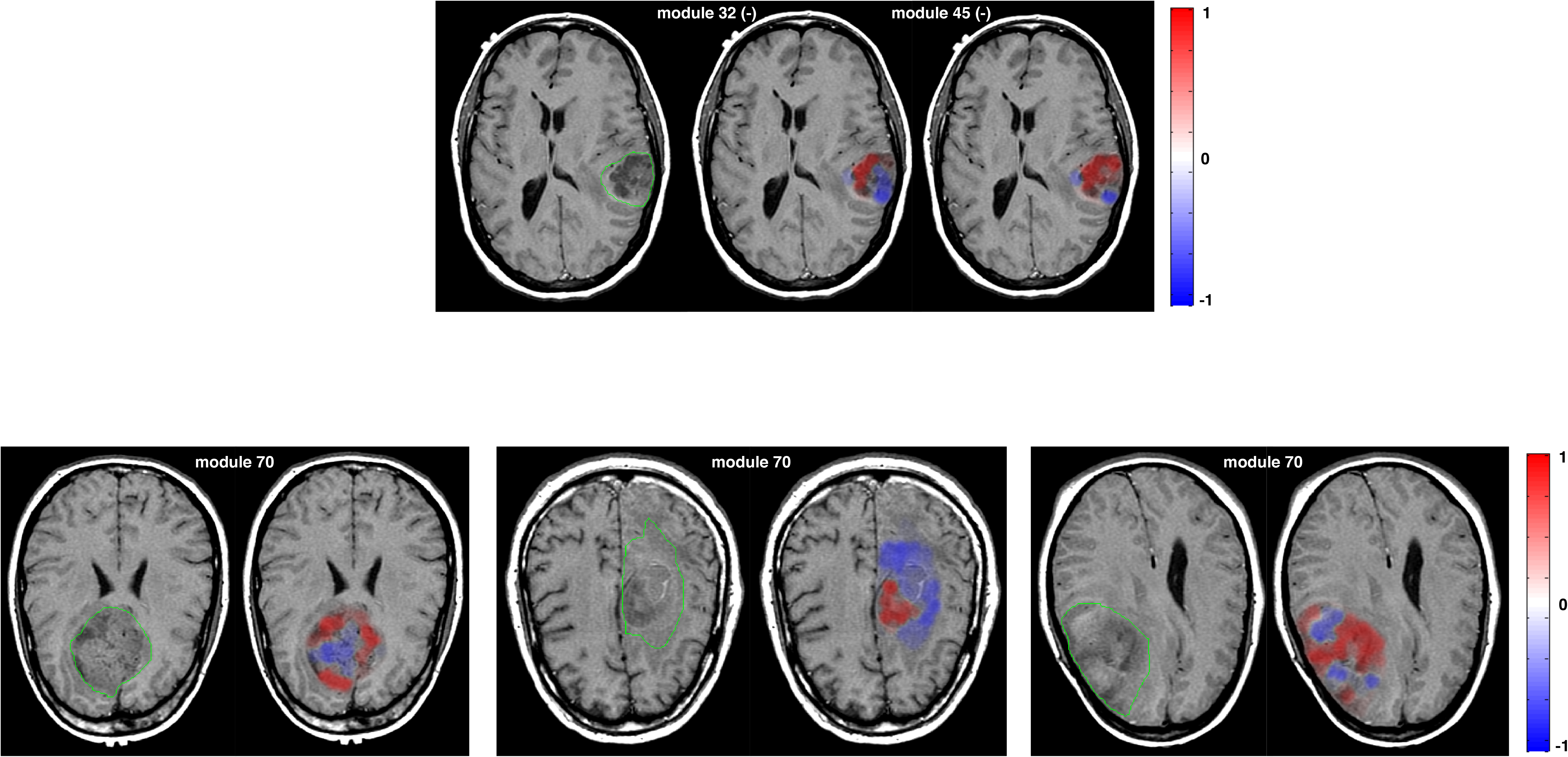
Comparison of expression heatmaps for modules 32 and 45 in the same patient, where both modules were globally under-expressed in the whole tumor. The axial views suggest that the under-expression does not come from consistently low MR signal for module 32, whereas only consistently high MR signal relates to the under-expression of module 45. This patient belongs to the REMBRANDT data and the heatmaps was generated using the model built on TCGA.

**Figure 4.**
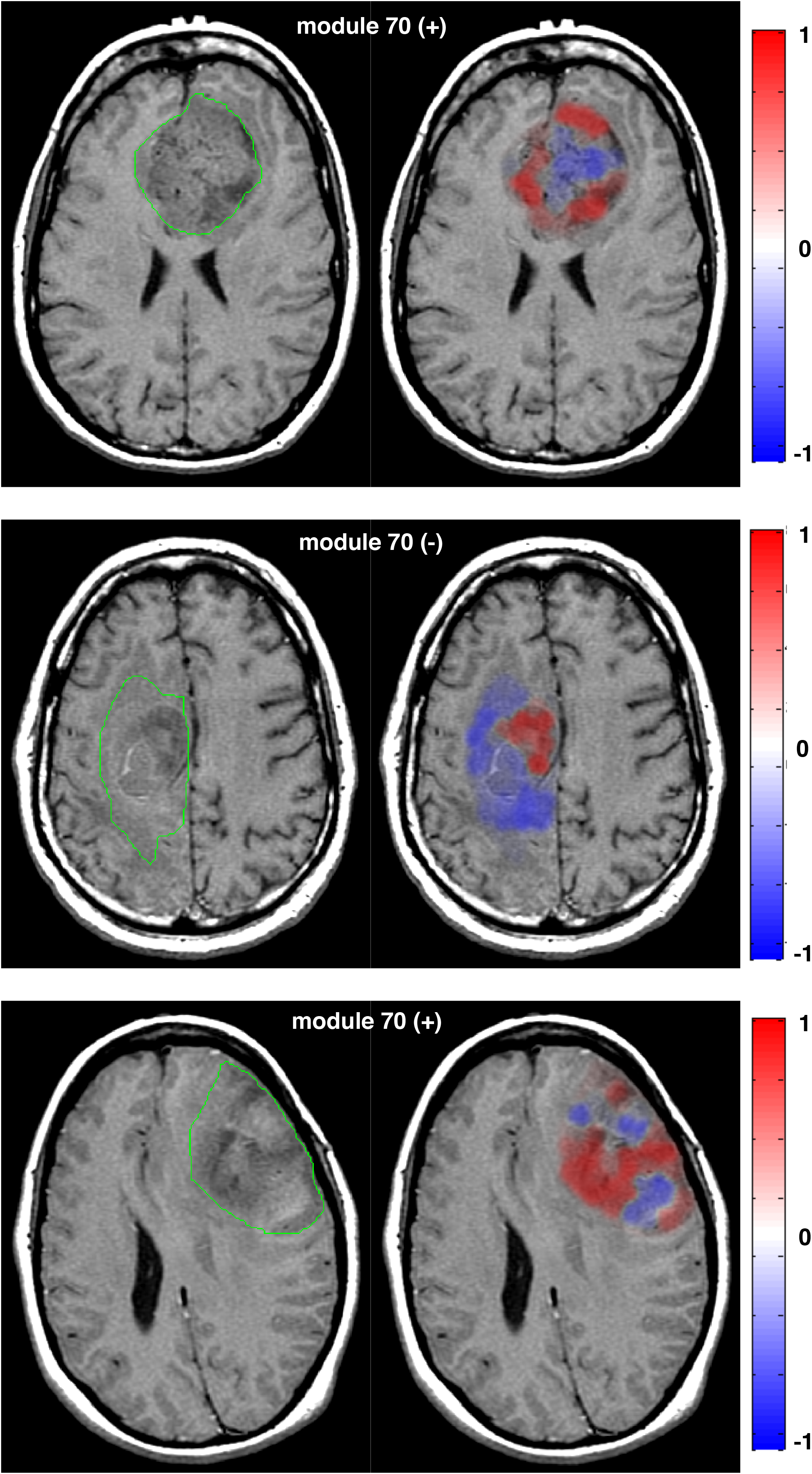
Expression heatmaps for module 70 in three different REMBRANDT patients when using the model built on TCGA. Its global expression was positive, negative and positive for the patients shown on top, middle and bottom, respsectively.Its over-expression is characterized by a consistent intermediate/low MR signal across patients.

### Validation of expression heatmaps

Because our measure of gene module expression is global over the tumor, the performance of patch-based local predictions of gene modules is biased towards morphologically uniform tumors (Table 1). We can, however, study the relation between predictive performance and the thresholds on the decision values (i.e., the output of the GLM model as defined in Eq. 1) to avoid forcing the model to provide a decision when patches have a weak absolute decision value. This shows that the AUC as a function of the threshold on the absolute decision values monotonously increases as a function of the predictive performance for most modules (Figure 5), suggesting that the decision values of the proposed models allow for a true identification of the local patches that are involved in the gene module expression. This indirectly validates the association between locoregional image patches and gene expression (Figures 3 and 4).

**Figure 5.**
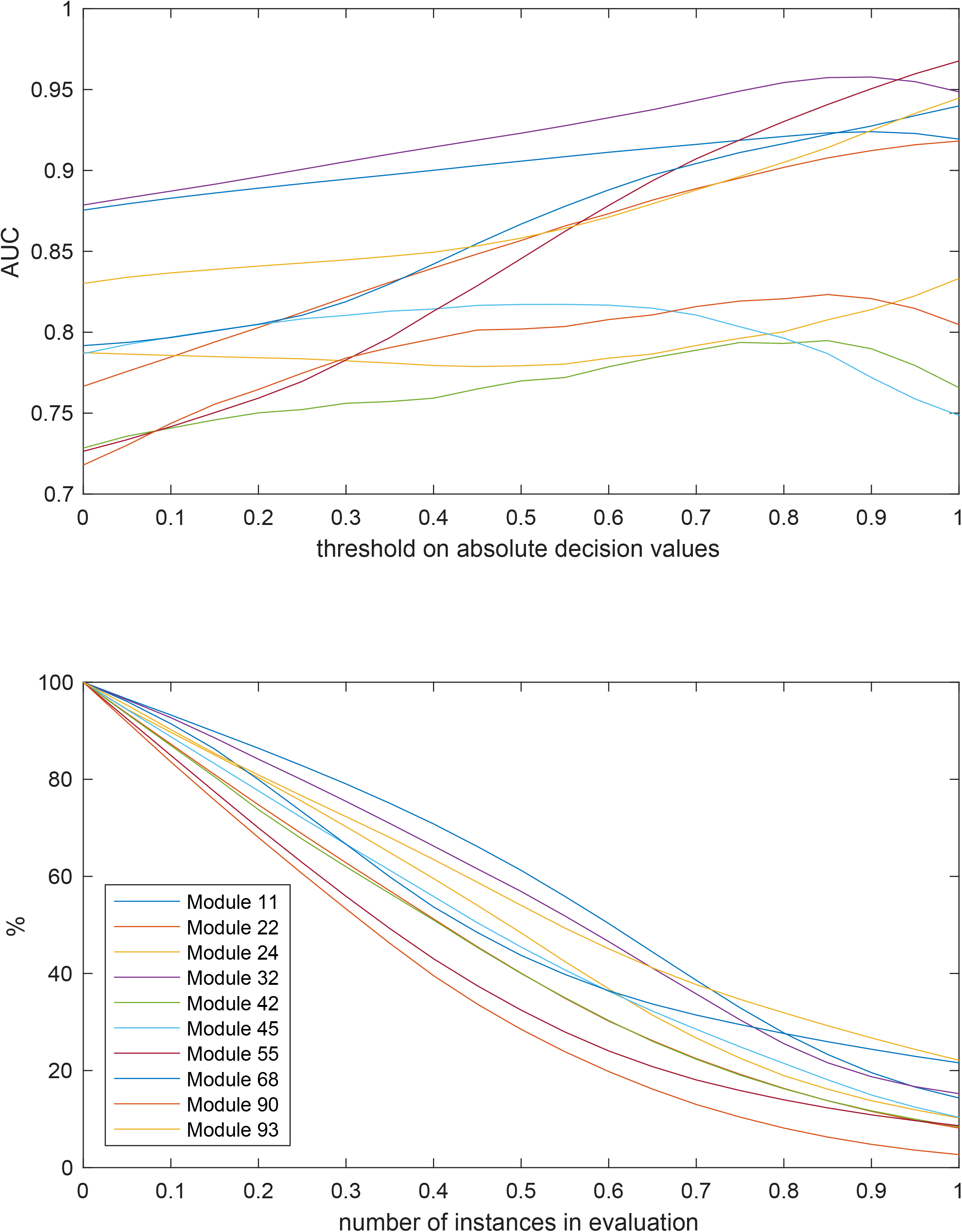
Evolution of AUC with respect to the threshold on absolute decision values (TCGA LOPO) for the top 10 gene modules shown in Table 1. The observed performance increase suggests that the model is able to truly identify the instances that are involved in the gene module expression.

## Discussion

We proposed a radiogenomics approach to study the intricate relationships between imaging phenotypes of GBMs and their molecular properties. Our approach relies on three main steps, including: (i) multi-omics data fusion to generate interpretable gene modules, (ii) locoregional image analysis and supervised multivariate models to establish multivariate models between MR image features and module expressions and (iii) the construction of module expression heatmaps to visualize intratumoral molecular heterogeneity as image overlays. When compared to previous works on radiogenomics of GBMs, we used automated image analysis with a finer grained level to take into account the locoregional heterogeneity of tumors and their molecular profiles. Secondly, the multi-omics clustering algorithm AMARETTO applied to the TCGA data allowed us to reduce data dimensionality while generating interpretable gene modules (Figure 2). Two thirds of the modules were also consistently observed on the validation cohort (REMBRANDT) with high R-square and module homogeneity (Supplementary Table 2). The proposed locoregional radiomics modeling was found to yield predictive AUCs on both cohorts for 15 modules (Table 1). For five of them, the model trained on TCGA could be used directly to predict their expression in REMBRANDT. For the other seven, significant AUCs were observed for both TCGA and REMBRANDT LOPO, but not when directly applying the model from TCGA on REMBRANDT. The number of features used in the models was at most 20 (i.e., the normalized intensity histogram bins), whereas the number of instances (i.e., patches) was in the range of 8,000 to 20,000. This ensured a number of instances far superior to the dimensionality of the model, thus limiting drastically the risk of overfitting as well as the risk of fortuitous discoveries and associations.

Next, we were able to associate biologically meaningful and clinically relevant interpretations for the best predicted gene modules. In particular, module 32 is involved in the maintenance of glioma stem cells, module 11 is a core cell cycle module and module 93 is involved in the receptor tyrosine kinase pathways. Finally, module 84 is linked to BCAN, a highly expressed gene in gliomas promoting the growth and motility of brain tumor cells. It is therefore meaningful to observe that module 84 is best predicted using patches extracted from the boundaries of the tumors and that BCAN expression can be deduced from MR imaging. Moreover, it has not escaped our attention that two drug targets are among the regulators of modules that can be predicted from imaging: TNFRSF1A and ERBB3. TNFRSF1A is a target of VB-111 an antiangiogenic agent that showed effectiveness in phase 1 trials ^51^ and is currently evaluated in a phase 3 trial (NCT02511405). Secondly, ERBB3 is part of the EGFR family, and an emerging cancer drug target^52^. Previous results have shown that targeting EGFR and EGFRvIII in glioblastoma results in compensatory mechanisms in other family members including ERBB3, therefore targeting ERBB3 is potentially equally important ^53,54^.

Texture features were not found to carry relevant information related to local heterogeneity (Supplementary Tables 3). While the latter is contradictory to the findings of ^15^ and ^16^ showing that texture attributes are correlated with the genotype of the tumor, a fundamental difference in our approach is the use of very localized image instances as patches with a radius of 12 pixels (6mm). It is difficult to extract rich texture information on these small ROIs, where the number of possible digital spatial frequencies is small and the influence of neighboring objects (e.g., skull or cortical gyri) is dominant^55^. Moreover, these previous reports used other texture feature types (i.e. Laws, Gabor, Haralick, local binary patterns and the discrete orthonormal Stockwell transform), which can also account the discrepant findings ^15^ and ^16^.

Next, the exploratory expression heatmaps are the first attempts to reveal the relationship between cancer driver gene module expressions and local imaging patterns using only global gene expression patterns. They provide the opportunity to identify imaging patterns on contrast-enhanced T1-weighted images that are most related to the expression of a given module (Figure 3 and 4). When used for gene modules that obtained both high predictive performance and biological interpretation, the proposed visualization opens up avenues for assessing and quantifying personalized response to treatment targeting precise genes by observing the presence and heterogeneity of the radiogenomics map.

Finally, we studied the relationship between thresholded GLM’s absolute decision values and the predictive performance to validate the predictions and the associated heatmaps. This avoided forcing models to provide a decision when the patches have low absolute decision values and showed that with increasing threshold value, the model performance consistently increased (Figure 5). This suggests that the proposed expression heatmaps are truly highlighting regions with high under- or over-expression of the modules. Combing our work with parallel efforts focusing on multiple sampling points (e.g., guided by the proposed expression heatmaps) in order to obtain local gene expression for validation promises to provide additional validation of the locoregional MR imaging patterns reflecting molecular properties of GBM ^16^. Taken together our radiogenomics framework provides future research direction for treatment allocation, treatment follow-up and treatment monitoring by using MR imaging phenotypes that reflect molecular properties of brain tumors. To conclude, we extended radiogenomics mapping to a locoregional level that can account for the molecular heterogeneity of GBM tumors. In addition to the demonstration of the feasibility of the latter with an external validation set, we built a radiogenomic map for clinically relevant gene modules with illustrative support from molecular expression heatmaps superimposed on medical images.

## Acknowledgements

Research reported in this publication was supported by the National Institute of Biomedical Imaging and Bioengineering of the National Institutes of Health under Award Number R01 EB020527. It was also supported by the Swiss National Science Foundation with grant agreements PZ00P2_154891 and 205320_179069. The content is solely the responsibility of the authors and does not necessarily represent the official views of the National Institutes of Health.

## Competing Interests

The authors declare that they have no competing financial interests.

## Correspondence

Correspondence and requests for materials should be addressed to O.G.

